# Deficiency of the lipid flippase ATP10A causes diet-induced dyslipidemia in female mice

**DOI:** 10.1101/2023.06.16.545392

**Authors:** Adriana C. Norris, Eugenia M. Yazlovitskaya, Lin Zhu, Bailey S. Rose, Jody C. May, Katherine N. Gibson-Corley, John A. McLean, John M. Stafford, Todd R. Graham

**Author notes:** Address correspondence to: Todd R. Graham, Department of Biological Sciences, Vanderbilt University, 465 21^st^ Ave S, Nashville, Tennessee 37212, USA.

## Abstract

Genetic association studies have linked ATP10A and closely related type IV P-type ATPases (P4-ATPases) to insulin resistance and vascular complications, such as atherosclerosis. ATP10A translocates phosphatidylcholine and glucosylceramide across cell membranes, and these lipids or their metabolites play important roles in signal transduction pathways regulating metabolism. However, the influence of ATP10A on lipid metabolism in mice has not been explored. Here, we generated gene-specific *Atp10A* knockout mice and show that *Atp10A*^-/-^ mice fed a high-fat diet did not gain excess weight relative to wild-type littermates. However, *Atp10A^-/-^*mice displayed female-specific dyslipidemia characterized by elevated plasma triglycerides, free fatty acids and cholesterol, as well as altered VLDL and HDL properties. We also observed increased circulating levels of several sphingolipid species along with reduced levels of eicosanoids and bile acids. The *Atp10A^-/-^* mice also displayed hepatic insulin resistance without perturbations to whole-body glucose homeostasis. Thus, ATP10A has a sex-specific role in regulating plasma lipid composition and maintaining hepatic liver insulin sensitivity in mice.

## Introduction

The metabolic syndrome is a complex condition that affects 33% of Americans (1) and on a global scale, over 1 billion people (2). This syndrome is characterized by abdominal obesity, insulin resistance, hypertension, and dyslipidemia, and an increased risk of developing type 2 diabetes mellitus and atherosclerotic cardiovascular disease (ASCVD) (3). Commonly found variants in *ATP10A* have been linked to increased risk of insulin resistance (4) and variants within a related gene, *ATP10D*, have been linked to increased atherosclerosis risk (5). ATP10A and ATP10D are P4-ATPases, also known as lipid flippases, that translocate lipids from the exoplasmic or luminal leaflets to cytosolic leaflets of cell membranes. This creates an asymmetric distribution of lipids within membranes that has several implications for the cell (6), including in apoptosis (7), vesicular trafficking (8), and signal transduction (9).

Interestingly, most of the 14 human P4-ATPases have established roles in disease, such as in severe neurological disorders and intrahepatic cholestasis (10). These enzymes transport specific lipid substrates, which are closely related to their cellular and physiological functions. ATP10A flips phosphatidylcholine (PC) (11) and glucosylceramide (GlcCer) (12). PC is the most abundant phospholipid in cellular membranes (13). Metabolites of PC, such as lysophosphatidylcholine (LysoPC), arachidonic acid (AA), and eicosanoids produced from AA, play important roles in modulating inflammation and various disease states (14–18). GlcCer is a bioactive lipid that can affect insulin signaling and energy homeostasis (19), as well as inflammation (20). Inhibiting GlcCer synthase enhances insulin sensitivity (21) and protects against cardiac hypertrophy (22). GlcCer can be broken down into ceramide or built up into gangliosides, which also have established roles in metabolism (23–25).

Previous reports suggested a role for murine ATP10A in diet-induced obesity, insulin resistance, and dyslipidemia and these phenotypes were more severe in female mice (26,27). However, the mouse model used in these studies contained a large, irradiation-induced chromosomal deletion (*p^23DFiOD^*) encompassing the pink-eyed dilution (*p)* locus, *Atp10A* (previously named *Atp10C)* and several other genes. End-point mapping of a nested deletion series implicated *Atp10A* in the metabolic defects but whether an *Atp10A*-specific knockout is sufficient to cause insulin resistance has not been determined. Homozygosity of the irradiation-induced deletion resulted in embryonic lethality and heterozygous mice that inherited the chromosomal deletion maternally displayed more severe metabolic phenotypes compared to those who inherited the deletion paternally. This observation suggested that the *Atp10A* locus was imprinted to suppress expression of the paternal allele, however several studies failed to detect imprinted expression of the *ATP10A* gene (28–30). It appears that *ATP10A* expression pattern is complex and dependent on multiple factors including gender, genetic background, and tissue type.

Although correlations between *ATP10A* deficiency and insulin resistance, diet-induced obesity, and glucose uptake (31) have been reported, it is unclear whether this flippase plays a causative role in these processes. Here, we generated a gene-specific deletion and tested the impact of *Atp10A* deficiency in mice during high fat diet (HFD) feeding. *Atp10A* deficient mice display female-specific diet-induced dyslipidemia, broad changes to lipid metabolism, and disruptions in liver insulin signaling.

## Results

### Atp10A deficiency does not affect body composition after high fat diet feeding

To develop the *Atp10A^-/-^* mice, we created a gene-specific knockout allele of *Atp10A* in the C57BL/6J background using CRISPR/Cas9 guide RNA sequences targeted to regions flanking exon 2 (Figure 1a) and confirmed the genotype of the mice via PCR (Figure 1b). To test the potential impact of *Atp10A* on weight gain, we placed male and female WT (*10A^+/+^*) and KO (*10A^-/-^*) mice on a HFD for 12 weeks. We found that there was no significant difference in weight gain in males (P-value for 12^th^ week on HFD: 0.3154) or females (P-value for 12^th^ week on HFD: >0.9999) based on genotype (Figure 1c). We next examined body composition and size and found that male and female mice did not display a difference in % lean mass (Figure 1d) or % fat mass (Figure 1e) based on genotype. Male mice also did not exhibit changes in body length based on genotype, however, female *Atp10A^-/-^* mice had shorter body lengths compared to *Atp10A^+/+^* littermates (Figure 1f). For the female mice, no significant difference was observed in daily (sum of light and dark hours) activity, food intake, or energy expenditure based on genotype. However, the *Atp10A^-/-^* female mice displayed reduced food intake and a negative energy balance during light hours compared to WT mice (Supplemental table 1). Additionally, we did not observe any changes in tissue mass (Figure 1g) in female mice based on genotype. We also measured weight gain, body composition, and body length in female mice on normal chow; and found no significant differences based on genotype (Supplemental Figure 1a-d). Altogether, these results indicate that *Atp10A* deficiency does not alter the development of diet-induced obesity under the conditions tested.

**Figure 1.**
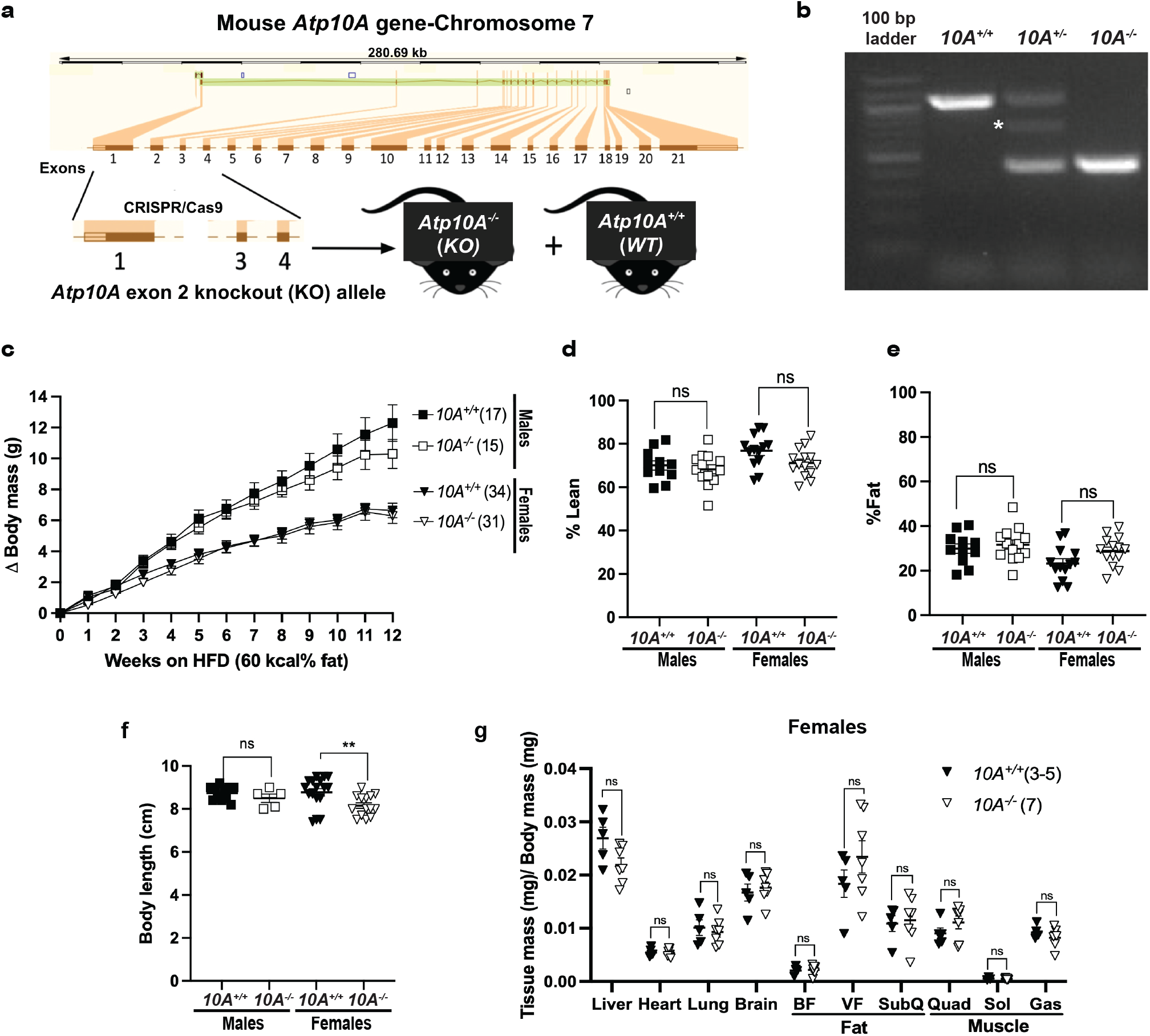
*Atp10A* deletion does not alter the development of diet-induced obesity after 12 weeks of HFD. (a) Graphic illustrating how CRISPR/cas9 was used to delete exon 2 in the mouse *Atp10A* to create the *Atp10A^-/-^* (KO) mouse line. (b) PCR genotyping results from *Atp10A^+/+^ (10A^+/+^*) (1067 base pairs (bp))*, Atp10A^+/-^* (*10A^+/-^*), and *Atp10A^-/-^ (10A^-/-^*) (460 bp) mouse tail clips. The white asterisk indicates a product at ∼700 bp that appears in all *10A^+/-^* samples due to hybridization of sequences from the *10A^+/+^* and *10A^-/-^* PCR products. (c) Weight gain of male and female *Atp10A^+/+^ (10A^+/+^*) and *Atp10A^-/-^ (10A^-/-^*) mice over the course of 12 weeks on a HFD (60 kcal% fat, Ad lib feeding), (**Male:** *10A^+/+^*n=17, *10A^-/-^* n=15; **Female:** *10A^+/+^*n=34, *10A^-/-^* n=31). (d) Lean and (e) fat body mass were normalized to the combined sum of lean and fat mass to calculate % Lean and % Fat mass (**Male**: *10A^+/+^* n=11,*10A^-/-^* n=13; **Female**:*10A^+/+^* n=13,*10A^-/-^* n=14). (f) Body length of mice was measured after CO_2_ sacrifice, **P=0.0094. (**Male:** *10A^+/+^* n=11, *10A^-/-^* n=5; **Female**: *10A^+/+^* n=17, *10A^-/-^* n=14). (g)The wet mass of tissue was measured after removal from female mice on the 12th week on the HFD. Tissue mass is normalized to body mass (*10A^+/+^* n=3-5,*10A^-/-^* n=7). P value by (a) 2-way ANOVA with Sidak’s multiple comparison or (d-g) unpaired t-tests.

### Atp10A deficiency does not affect whole-body glucose homeostasis in female mice

Studies using the *p^23DFiOD^* deletion encompassing *Atp10A,* found that these mice were hyperinsulinemic, insulin resistant, and hyperglycemic (26,27). To specifically assess the influence of *Atp10A* on glucose homeostasis, we fasted WT and *Atp10A^-/-^* (exon 2 deletion) mice for 5 hours and performed an oral glucose tolerance test (OGTT) as well as measured fasting levels of glucose and insulin. During the OGTT, male *Atp10A* deficient mice did not exhibit differences in glucose excursion after the glucose bolus (Figure 2a) compared to WT mice. However, the female *Atp10A*^-/-^ mice exhibited significantly elevated fasting blood glucose after a 5 hour fast (points before time 0 during the OGTT, Figure 2b) and trended toward exhibiting an elevated glucose excursion compared to the female WT mice, represented by the elevated area under the curve (AUC) (Figure 2b, inset). While the elevated fasting blood glucose was significant for the number of female mutant mice tested (n=6), the study was underpowered. Therefore, fasting blood glucose was measured from additional mice and we found no significant difference in fasting blood glucose in male or female mice based on genotype (Figure 2c). The reason for the variation in fasting blood glucose seen in the females is unclear, but could be due to experimental stress for the animals with the elevated fasting blood glucose There was also no significant difference in fasting plasma insulin (Figure 2d) or insulin levels at 30 minutes (T30, Figure 2e), or 60 minutes (T60, Figure 2f) after the glucose bolus (during the OGTT) in males or females based on genotype. Additionally, normal chow fed female *Atp10A^-/-^* mice did not exhibit differences in fasting blood glucose or insulin (Supplemental Figure 1e,f) compared to WT mice. Therefore, in contrast to expectations based on prior reports, *Atp10A* deficiency does not appear to significantly perturb whole-body glucose homeostasis.

**Figure 2.**
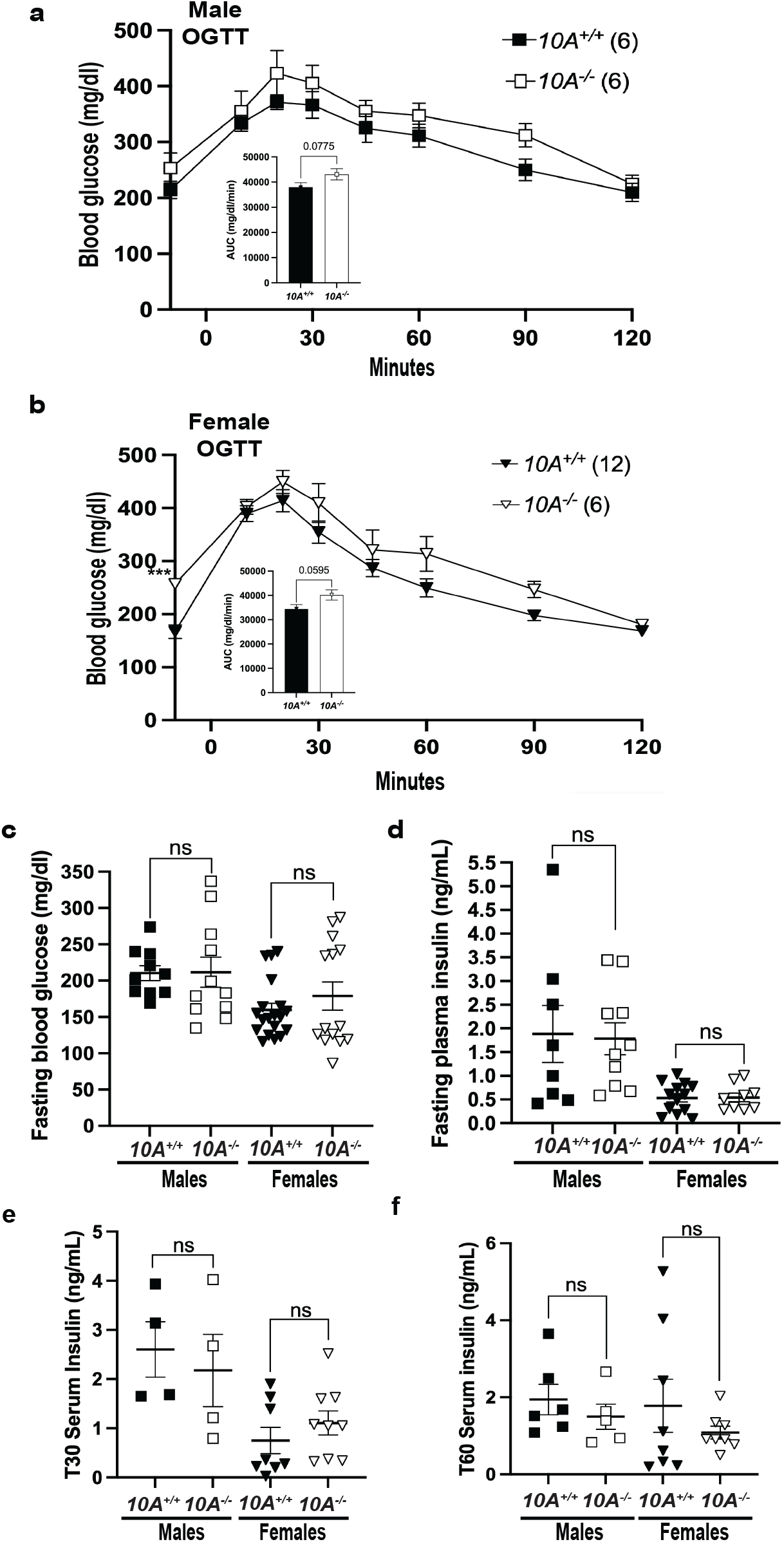
*Atp10A* deletion does not result in changes to glucose homeostasis after 12 weeks of HFD. **(a)** Male and (b) female mice were gavaged with 20% dextrose (time 0 on the graph) after a 5-hour fast then an oral glucose tolerance test (OGTT) was performed. The inset shows the area under the curve (AUC) of the blood glucose (mg/dl) during the OGTT, normalized to fasting blood glucose (points before time 0 on the graph), ***P=0.0003. **Male:** *10A^+/+^* n=6,*10A^-/-^* n=6; **Female**: *10A^+/+^* n=12, *10A^-/-^* n=6) .(c) Fasting blood glucose was measured from tail blood after a 5 hour fast, via a glucometer **(Male:** *10A^+/+^* n=10, *10A^-/-^* n=11; **Female**: *10A^+/+^* n=19, *10A^-/-^* n=14). (d) Fasting plasma insulin was measured after a 5 hour fast (**Male:** *10A^+/+^* n=8, *10A^-/-^* n=10; **Female:** *10A^+/+^* n=13,*10A^-/-^* n=9). Insulin was measured from serum collected during the OGTT shown in Figure 2a,b at the (e) 30-minute time point (T30) and the (f) 60-minute time point (T60), (**Male:** C: *10A^+/+^* n=4, *10A^-/-^* n=4, D: *10A^+/+^* n=6, *10A^-/-^* n=5; **Female**: *10A^+/+^* n=8, *10A^-/-^* n=6). P value by (a,b, not AUC) 2-way ANOVA with Sidak’s multiple comparison or (a,b (AUC), c-f) unpaired t-tests.

### Maternal inheritance of the Atp10A exon 2 deletion results in changes to body length and fasting blood glucose in males

Prior reports showed that the inheritance pattern of the *p^23DFiOD^* deletion encompassing *Atp10A* effected the severity of the metabolic phenotypes in heterozygous mice, specifically if the deletion was inherited maternally or paternally (26,27). We tested this phenomenon using the *Atp10A* exon 2 deletion mice by measuring metabolic phenotypes after high fat diet feeding, in heterozygous mice reared from *10A^+/-^* dams x *10A^+/+^* sires (maternal inheritance (inh.)) or *10A^+/+^* dams x *10A^+/-^* sires (paternal inh.). We found no significant difference in weight gain, body composition, or fasting plasma insulin based on the inheritance patterns of the *Atp10A* exon 2 deletion in males or females after 12 weeks of HFD (Supplemental Figure 2a-c, f). However, the male mice inheriting the exon 2 deletion maternally trended toward having increased weight gain, although this did not reach statistical significance (P-value for 12^th^ week of HFD: 0.0906), shorter bodies (Supplemental Figure 2d) and elevated fasting blood glucose levels after a 5 hour fast (Supplemental Figure 2e) compared to mice that inherited the deletion paternally. Therefore, the inheritance pattern of the *Atp10A* exon 2 deletion does not significantly impact weight gain, body composition, and fasting plasma insulin in males or females after HFD feeding, but does have distinct effects on body length and fasting blood glucose levels in male mice.

### Atp10A deficiency causes diet-induced dyslipidemia in females

To further probe the influence of *Atp10A* on whole-body metabolism, we measured the concentration of free fatty acids (FFA), cholesterol (chol), and triglycerides (TG) in the plasma. While the male mice showed no difference in plasma lipids based on genotype (Supplemental Figure 3a-c), the female *Atp10A^-/-^* mice display substantially elevated plasma concentrations of FFA (Figure 3a), chol (Figure 3b), and TG (Figure 3c) compared to *Atp10A^+/+^* littermates. To further explore this female-specific dyslipidemia phenotype, we performed fast performance liquid chromatography (FPLC) on plasma to assess the relative size and lipid distribution in lipoprotein fractions (Figure 3d). We found that the *Atp10A^-/-^*mice carry most of their TG and cholesterol in smaller sized very low-density lipoprotein (VLDL) and high-density lipoprotein (HDL) particles, respectively (indicated by a shift to the right in the chromatograms). We also compared the AUC of the VLDL-TG (Figure 3e) and HDL-cholesterol (Figure 3f) fractions and found that the *Atp10A^-/-^* mice trend toward having more lipids in these fractions compared to controls, although this did not reach statistical significance. Together, these results show that *Atp10A* deficiency causes female-specific dyslipidemia and changes to lipoprotein metabolism after HFD feeding.

**Figure 3.**
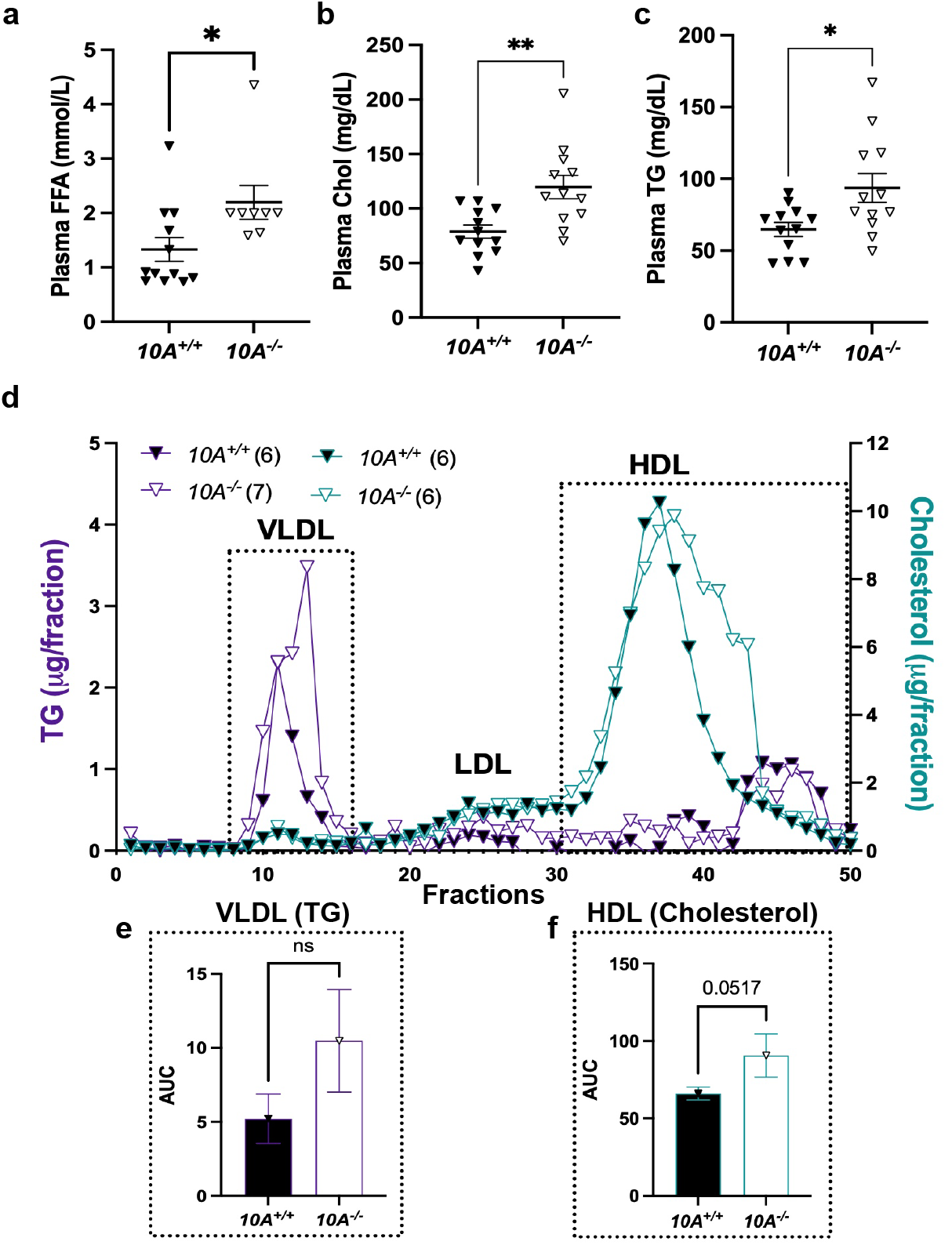
*Atp10A* deletion causes diet-induced dyslipidemia in female mice after 12 weeks of HFD. (a-c) Free fatty acids (FFA), cholesterol (chol), and triglycerides (TG) were measured in plasma from females after a 5 hour fast or after a 5 hour fast followed by an OGTT (a, *10A^+/+^*n=12, *10A^-/-^* n=9,*P=0.0320; b, *10A^+/+^* n=12, *10A^-/-^* n=9, **P=0.0030; c, *10A^+/+^* n=12, *10A^-/-^* n=9, *P=0.0165.) (d-f) Lipoproteins in pooled plasma from females, collected after a 5-hour fast or 5-hour fast and OGTT, were separated via FPLC. (e) Plasma TG (purple) and (f) cholesterol (blue) content were measured from the separated lipoproteins. AUC of (e) TG fractions 10-18 of VLDL and (f) cholesterol fractions 30-50 of HDL (**TG:***10A^+/+^* n=6, *10A^-/-^* n=7, **Chol:***10A^+/+^* n=6, *10A^-/-^* n=6). P value by unpaired t-test.

### Atp10A deficiency in females causes substantial changes to the plasma lipidome and visceral fat transcriptome after HFD feeding

To get a broader view of the dyslipidemia in the plasma of female *Atp10A^-/-^* mice, we performed mass spectrometry-based untargeted lipidomics on plasma from female *Atp10A^-/-^*and *Atp10A^+/+^* littermates after 12 weeks on HFD after a 5-hour fast. We found that *Atp10A* deficiency resulted in statistically significant changes to the abundance of a large number of plasma lipid species; 591 in total, with 324 of these observed from positive ion mode and 267 from negative ion mode lipidomics (Figure 4a, Study ID ST002696, http://dx.doi.org/10.21228/M83H7N). To highlight how *Atp10A* deficiency altered specific classes of lipid species, we plotted the log2(Fold Change) of lipid species that were significantly changed and calculated the % of the lipid species in each group that had a positive fold change (increase in abundance, green values) versus a negative fold change (decrease in abundance, blue values) (Figure 4b). Interestingly, we saw modest increases in the abundance of ATP10A’s lipid substrates, PC and hexosylceramides (this includes GlcCer and galactosylceramide (GalCer)). Surprisingly, we observed that plasma eicosanoid, bile acid and fatty acid (FA) conjugate species were dramatically depleted in *Atp10A^-/-^* mice compared to *Atp10A^+/+^* littermates.

**Figure 4.**
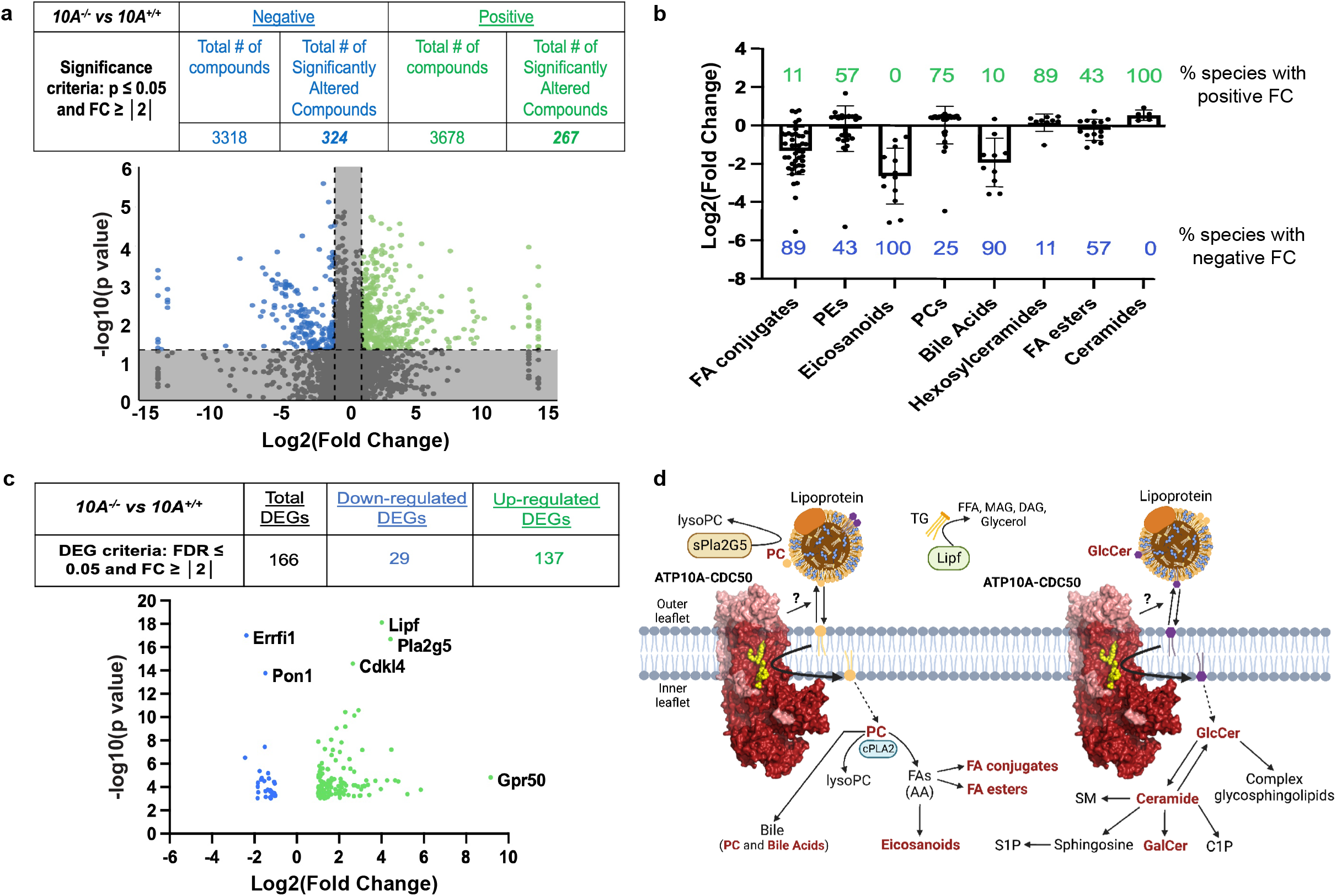
*Atp10A* deletion leads to substantial perturbations in the plasma lipidome and visceral fat transcriptome in female mice on a HFD. (a) Lipid metabolites from fasting plasma were measured by HPLC-IM-MS/MS and the data was processed via Progenesis QI. The table indicates the number of total and significantly changed lipid metabolites (compounds) observed from each ionization mode (Negative and Positive); the volcano plot illustrates the fold changes (FC) of compounds between *10A^-/-^* vs. *10A^+/+^* mice, the grey shading indicates changes that did not meet the significance criteria. P-values by ANOVA. (*10A^+/+^* n=8, *10A^-/-^* n=8, see Supplemental Table 3 for complete information about sample size) (b) The log2(FoldChange) of lipid metabolites compared between *10A^-/-^* vs. *10A^+/+^* mice are grouped by indicated categories (green and blue indicate the % of lipid species with positive and negative fold change, respectively, between *10A^-/-^* vs. *10A^+/+^* mice; P values ≤ 0.1 by ANOVA). (c) Sequencing of visceral fat mRNA was performed using the Illumina NovaSeq 6000. The table provides the criteria used to calculate the number of differentially expressed genes (DEGs) and how many DEGs were measured; the volcano plot illustrates the DEGs (*10A^+/+^* n=3,*10A^-/-^* n=3). (d) Schematic illustrating the metabolic pathways linking ATP10A substrates to observed changes in the plasma lipidome and visceral fat transcriptome. Lipid metabolites with significant changes due to *10A* deletion from panel B are in bold red, the hexosylceramide category includes both GalCer and GlcCer. The protein products from two mRNA transcripts that were significantly upregulated in the visceral fat from *10A^-/-^*mice (panel c), Pla2G5 and Lipf, are highlighted to show their role in lipid metabolism. Schematic created using Biorender.com. See Supplemental Table 2 for RNASeq KEGG pathway analysis. PC= phosphatidylcholine, AA=arachidonic acid, TG= triglyceride, FA=fatty acids, FFA=free fatty acids, DAG=diacylglycerol, MAG=monoacylglycerol, SM=sphingomyelin, S1P= sphingosine 1-phosphate, C1P= ceramide 1-phosphate, GalCer= galactosylceramide, GlcCer= glucosylceramide.

We further probed the influence of *Atp10A* on metabolism by performing RNAseq on visceral fat (Figure 4c, NCBI GenBank accession numbers: SRR24233646, SRR24233645). We found that 166 genes were differentially expressed in *Atp10A^-/-^*compared to *Atp10A^+/+^* mice, where 29 were downregulated and 137 were upregulated. Some notable differences were substantial increases in expression of *Lipf* and *Pla2g5,* encoding a TG lipase and a secreted phospholipase A_2_, respectively, in *Atp10A^-/-^* mice (Figure 4c, Supplemental Table 2). Additionally, the orphan G protein-coupled receptor, GPR50, was upregulated 571-fold. This gene has been linked to circulating TG and HDL levels in humans (32), energy metabolism in mice (33), and the attenuation of inflammation and insulin signaling in 3T3-L1 adipocytes (34).

Taken together, these results indicate that ATP10A has a substantial role in lipid metabolism in female mice fed a HFD. The potential mechanistic links between the lipid transport activity of ATP10A and whole-body lipid metabolism are shown in Figure 4d. Briefly, by translocating PC and GlcCer from the outer leaflet to the inner leaflet of the plasma membrane, ATP10A provides substrates to intracellular lipid metabolism enzymes for production of bioactive lipid signaling molecules while decreasing the availability of PC and GlcCer to molecules in circulation (i.e. lipoproteins, lipases) (Figure 4d). Thus, this lipid transport could affect levels of PC, GlcCer, and downstream metabolites in the intracellular space and in circulation (Figure 4b) as well as change the expression levels of lipid handling enzymes (Figure 4c).

### Atp10A deficiency in females results in changes to liver lipid metabolism after HFD feeding

To further probe the impact of *Atp10A* deficiency on lipid metabolism, we measured levels of various lipid species in the liver. We found that female *Atp10A^-/-^* mice have less liver FFA (Figure 5a), but no significant differences in the levels of total cholesterol, cholesterol esters (CE), unesterified cholesterol, TG, phospholipids (PL), or ceramides compared to *Atp10A^+/+^* littermates (Supplemental Figure 4a-f). Female *Atp10A^-/-^* mice also exhibit a significant increase in the abundance of monounsaturated vs. saturated species of several TG and PL species (Figure 5b) and one CE species (Figure 5c) with carbon chain lengths of 16 and/or 18. Although the total amounts of cholesterol and TG in the liver were not different based on genotype, Oil Red O stained lipid droplets (LD) from the *Atp10A^-/-^* mice were larger on average compared to *Atp10A^+/+^* mice (Figure 5d-f). Therefore, *Atp10A* deficiency results in changes to the amount and saturation state of several lipid species in the liver.

**Figure 5.**
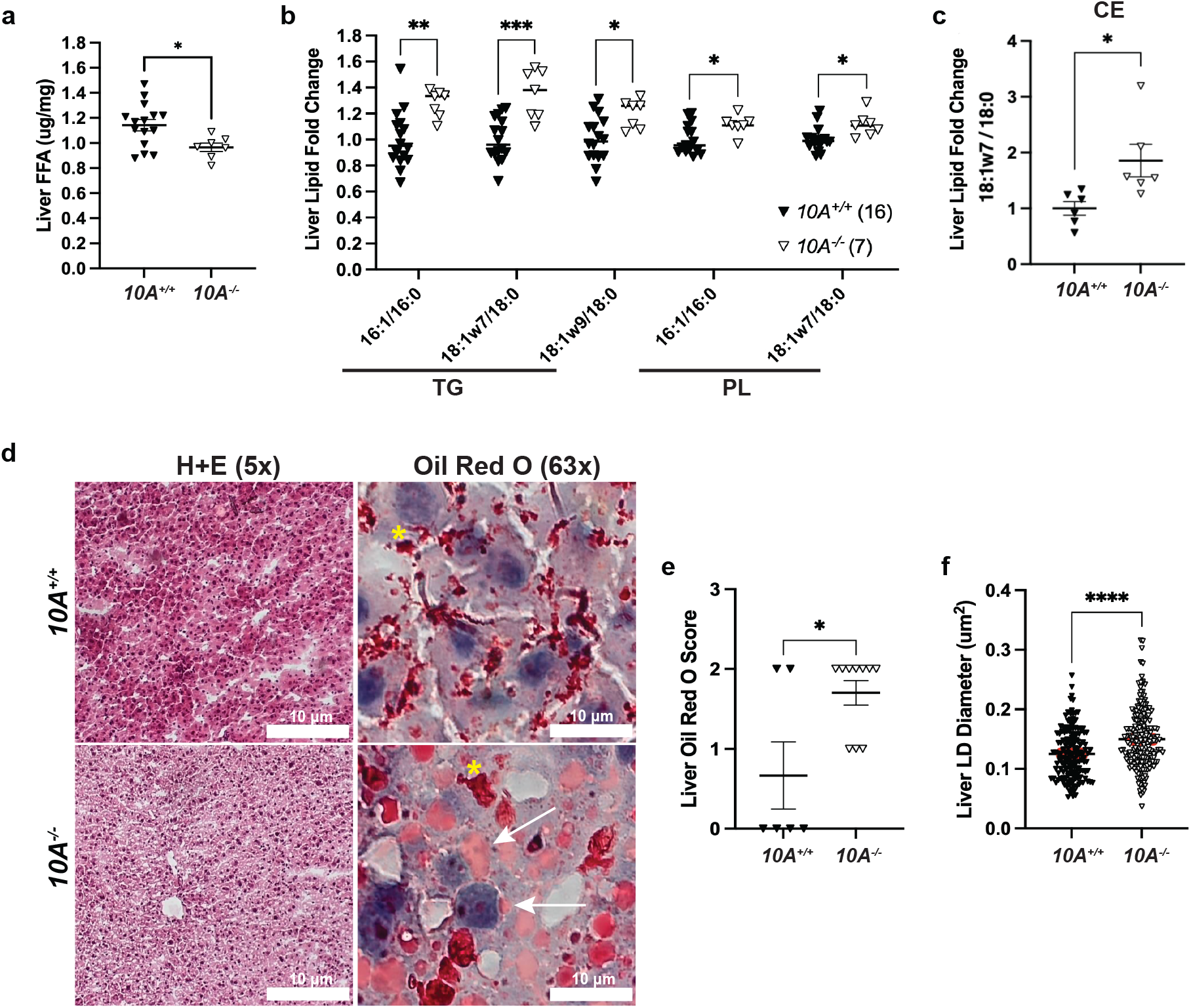
*Atp10A* deletion causes changes to liver lipid metabolism in female mice after 12 weeks of HFD. (a) FFAs were measured from flash frozen livers via gas chromatography. Livers were collected after a 5 hour fast or after a 5 hour fast followed by an OGTT. t (FFA: *10A^+/+^* n=15,*10A^-/-^* n=7, *P=0.0201). The saturation of liver (b) TG, PL, and (c) CE species was determined by gas chromatography. The fold change of monounsaturated vs saturated TG, PL, CE is shown, **TG**: **P=0.0025, ***P=0.0001, *P=0.0443, **PL**: *P=0.0297, *P=0.0281 (*10A^+/+^* n=16, *10A^-/-^* n=7). **CE:** *P=0.0223 (*10A^+/+^* n=6, *10A^-/-^* n=6). (d) Representative images of livers stained with H+E (5x) or Oil Red O (63x). The coalescence of the Oil Red O stain is indicated by yellow asterisks (artifact) and the arrows point to neutral lipids stained by Oil Red O. Scale bars = 10 μM. (e) Liver sections were scored using the Oil Red O Score described in the Materials and Methods, *P=0.0327 (*10A^+/+^* n=6, *10A^-/-^* n=10). (f) The diameters of Oil Red O positive lipid droplets (LD) in the liver sections with a score of 2 were measured using ImageJ. The red line in the graph represents the SEM, ****P=<0.0001 (*10A^+/+^*,120 LDs measured, n=2, *10A^-/-^*, 203 LDs measured, n=3). P value by (a-c, f) unpaired t-test or (e) Mann-Whitney U test.

### Atp10A deficiency results in changes to liver metabolic signaling

Next, we evaluated potential hepatic mechanisms responsible for the metabolic phenotypes observed in the HFD fed female mice by examining key regulators of energy and lipid metabolism in the liver (Figure 6f). Diacylglycerol acyltransferases (DGATs) catalyze the final step in TG synthesis (Figure 6f) and we found that DGAT2 was significantly elevated in liver from HFD-fed *Atp10A^-/-^* females (Figure 6a,b). These results indicate that the lower liver FFA and larger lipid droplets in the hepatocytes might be due to the higher activity of DGAT2 in *Atp10A^-/-^* mice. AMP kinase (AMPK) is a central regulator of energy metabolism that inhibits acetyl-CoA carboxylase (ACC) activity by phosphorylating it at S79 (Figure 6f). Western blot analysis showed that the phosphorylation of AMPKα at T172 and AMPK-dependent inhibitory phosphorylation of ACC at S79 were significantly increased in *Atp10A^-/-^* females (Figure 6a,c), which may further contribute to the decreased liver FFA in *Atp10A^-/-^* mice. Of note, insulin signaling including the activating phosphorylation of the insulin receptor β subunit (IRβ) at Y1146, insulin receptor substrate-1 (IRS-1) at S612, and Akt at S473/T308 were decreased in *Atp10A^-/-^* mice even though glucose tolerance was not significantly changed (Figures 6d,e and 2b). Consistent with the changes in insulin signaling, the Akt-dependent inhibitory phosphorylation at GSK-3β at S9 was also decreased in *Atp10A^-/-^* females (Figure 6d,e). Cytosolic phospholipase 2 (cPLA_2_) is an enzyme that catalyzes the hydrolysis of phospholipids, such as PC, to LysoPLs and AA, and AA can be oxidized to form eicosanoids (35). Interestingly, the activating phosphorylation of cPLA_2_ at S505 was decreased in the *Atp10A^-/-^* mice (Figure 6d,e), which might contribute to the decreases in eicosanoids in plasma from *Atp10A*^-/-^ mice (Figure 4b). Altogether, these studies reveal that *Atp10A* deficiency in females fed a HFD results in several alterations to metabolic signaling in the liver, including depression of the insulin signaling pathway.

**Figure 6.**
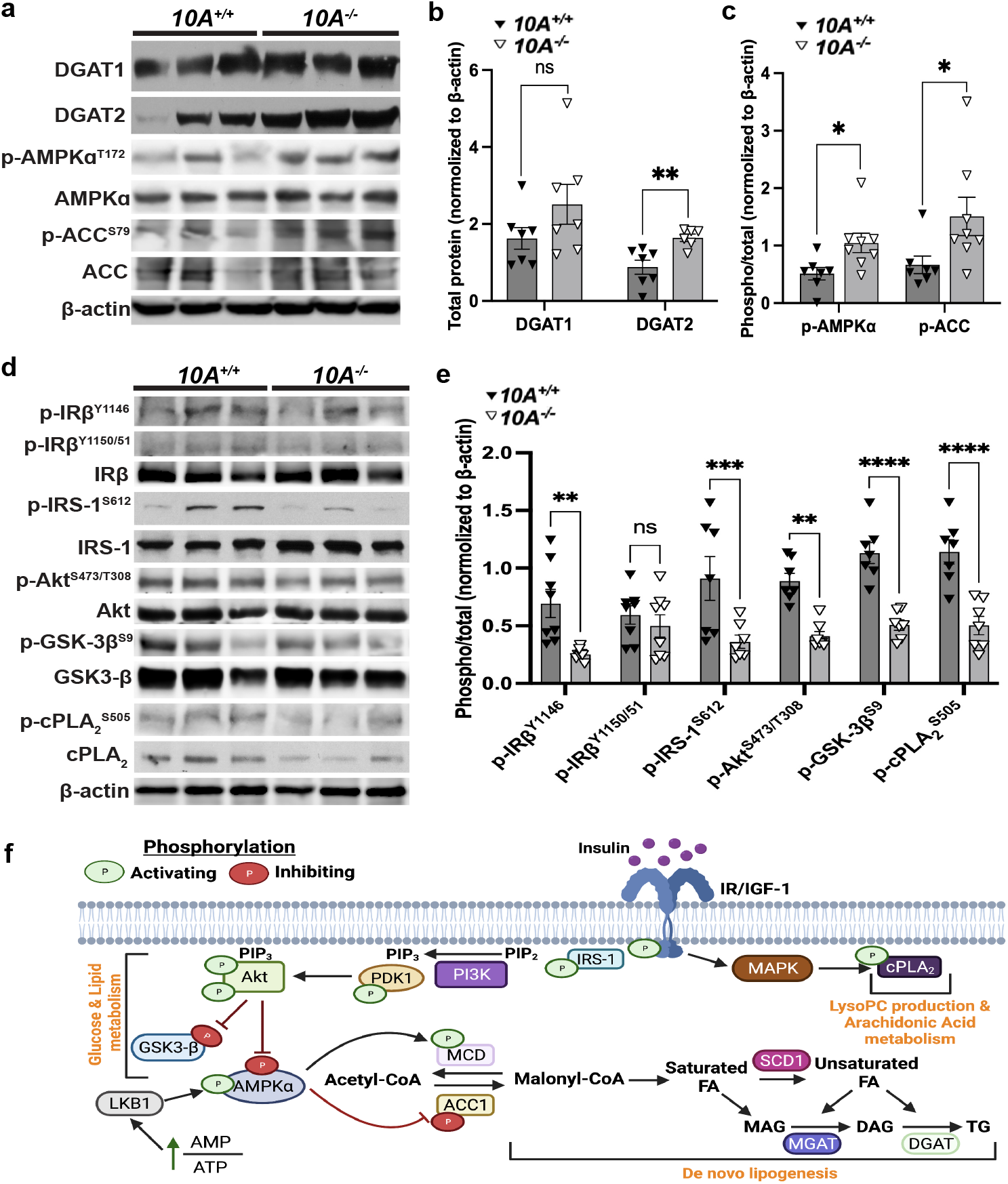
*Atp10A* deletion perturbs liver insulin signaling in female mice fed a HFD. (a, d) Representative blots for total and/or phosphorylated (a) DGAT1, DGAT2, AMPKα, ACC, and (d) IRβ, IRS-1, Akt, GSK-3β, and cPLA_2_. (b) For DGAT1 and DGAT2 quantitation, total protein was normalized to β-actin levels. (c, e) The phosphoproteins were normalized to their respective total protein and β-actin levels. Mean measurements of 4-6 independent experiments are shown. P value by (b, c) unpaired t-test or (e) 2-way ANOVA with Sidak’s multiple comparison test. (a-c, *10A^+/+^* n=7, *10A^-/-^* n=7, b: **P=0.0050; c: *P=0.0229, *P=0.0481; d, e, *10A^+/+^* n=8, *10A ^/-^* n=8, **P=0.0044, ***P=0.0007, **P=0.0038, ****P=<0.0001,). (f) Schematic adapted from (56) and created using Biorender.com. The illustration shows insulin binding to the insulin receptor and the resulting downstream signaling events that stimulate LysoPC production, AA metabolism, changes to glucose and lipid metabolism; including the promotion of lipogenesis. MAPK= mitogen-activated protein kinase, cPLA_2_= cytosolic phospholipase 2, IRS-1= insulin receptor substrate 1, PI3K= phosphoinositide 3-kinase, PIP2=phosphatidylinositol 4,5-bisphosphate, PIP3= phosphatidylinositol 3,4,5-triphosphate, PDK1= pyruvate dehydrogenase kinase 1, GSK-3β= glycogen synthase kinase-3 beta, LKB1= liver kinase B1, AMPKα= AMP-activated protein kinase alpha, MCD= malonyl-CoA decarboxylase, ACC1=acetyl-CoA carboxylase, SCD1= stearoyl-CoA desaturase 1, MGAT=monoglycerol acyltransferase, DGAT= diglyceride acyltransferase.

## Discussion

This study reveals a requirement for ATP10A in protecting female mice from dyslipidemia after HFD feeding. Despite the normal weight gain and body composition (Figure 1), female *Atp10A^-/-^*mice display elevated plasma levels of FFA, cholesterol, and TGs, and these parameters are unchanged in male *Atp10A^-/-^* mice relative to WT controls (Figure 3, Supplemental Figure 3). The plasma lipidome, visceral fat transcriptome, as well as hepatic lipid metabolism are also substantially perturbed in the knockout female mice (Figure 4,5). Additionally, basal insulin signaling in the liver is depressed (Figure 6) although the plasma insulin levels are comparable to wild-type littermates (Figure 2d).

*Atp10A* deficient female mice exhibit dyslipidemia characterized by elevated plasma FFA, cholesterol, TG, and alterations in the size and lipid content of both VLDL and HDL (Figure 3a-f). A potential source of hyperlipidemia is *de novo* lipogenesis in the liver; however, *Atp10A* deficient mice exhibit decreased total liver FFAs (Figure 5a) and an increase in activating phosphorylation of hepatic AMPKα (Figure 6a,c), a master regulator of lipid metabolism, that increases FA oxidation and decreases *de novo* lipogenesis through inhibition of ACC (36). Consistently, we also observed elevated inhibitory AMPKα-dependent phosphorylation of ACC (Figure 6a,c). Thus, it is unlikely that de novo FA production in the liver is driving the excess plasma TG and FFAs. It is possible that an increased release of FFA from dietary lipids or adipose tissue drives an increase in TG synthesis in the liver and its secretion in VLDL (Figure 3g) (37–39). Indeed, we observed an increase in DGAT2 expression in the liver, which could account for the elevated TG. In addition, *Lipf* mRNA transcripts were elevated in visceral fat from *Atp10A* deficient mice (Figure 4c, Supplemental Table 2). *Lipf* codes for a gastric lipase that breaks down TG into glycerol and FFAs and has been shown to be highly expressed in adipose tissue in a mouse model of diet-induced obesity (40). It is possible that *Atp10A* deficiency exacerbates this phenomenon, even though the WT and mutant mice exhibit no difference in weight gain or adiposity during high fat diet feeding, via an unknown mechanism.

Loss of ATP10A also disrupts cholesterol metabolism in the female mice. Plasma cholesterol is elevated and the HDL particles carrying the cholesterol are skewed towards smaller sizes. Additionally, *Atp10A* deficient mice show reduced circulating levels of bile acids (Figure 4a,b), which are important signaling lipids with implications in cholesterol metabolism and cardiovascular health (41). Further research is needed to understand the mechanisms underlying the perturbations to cholesterol metabolism observed in *Atp10A* deficient mice.

The molecular basis for the female-specific dyslipidemia in *Atp10A*^-/-^ mice is unclear. There are established differences in disease severity and prevalence between men and women (42). Premenopausal women tend to have lower cardiovascular risks compared to men and there are both hormonal and genetic contributions to sexual dimorphisms in lipid metabolism (43,44). The *Atp10A* promoter is predicted to have a transcription factor binding site for estrogen receptor alpha (Genecards.org, QIAGEN), so it is possible that differences in expression due to sex contribute to the consequences of *Atp10A* deletion. It is also possible that sex hormones either suppress the effect of *Atp10A^-/-^* on lipid metabolism in males or promotes these phenotypes in females. We did not measure the lipoprotein profiles, perform plasma lipidomics, or measure metabolic liver signaling pathways in male mice because they did not exhibit diet-induced dyslipidemia (Supplemental Figure 3a-c). Interestingly, we found that WT male mice exhibited higher levels of fasting FFA and cholesterol than WT females, suggesting a sex-based difference in the kinetics of lipid clearance during the postprandial response (Figure 3a-c, Supplemental Figure 3a-c). There is a need for a comprehensive investigation comparing fasting levels of lipids between male and female mice after HFD feeding to uncover potential sexual dimorphisms and elucidate the underlying mechanisms driving such differences.

*Atp10A* deficient female mice display elevated levels of monounsaturated versus saturated TG, PL, and cholesterol ester (CE) species in their livers (Figure 5b,c). This could be due to an increase in activity of stearoyl-CoA desaturase–1 (SCD1), an enzyme that catalyzes the synthesis of monounsaturated FAs from saturated FAs (Figure 6f). Interestingly, Scd1 activity has been suggested to be required for the onset of diet-induced hepatic insulin resistance (45). Liver sections from *Atp10A* deficient mice exhibit increased Oil Red O staining, a stain that binds neutral lipids, as well as larger lipid drops compared to control mice (Figure 5d-f). We observed an increase in the expression of DGAT2 (Figure 6b) in the liver of *Atp10A* deficient mice, but no significant increase in liver TG (Supplemental Figure 4d) or cholesterol levels (Supplemental Figure 4a-c), therefore the increase in Oil Red O staining could be due to staining of another neutral lipid and this needs to be further explored. Furthermore, female *Atp10A* deficient mice display impaired insulin signaling in the liver (Figure 6), highlighting the need for further exploration to elucidate the relationship between dyslipidemia, liver lipid metabolism, and altered insulin signaling observed in these mice.

Our studies suggest that *Atp10A* deficiency disrupts the regulation of cPLA_2_ activity and the production of eicosanoids. We found a reduction of activating phosphorylation of cPLA_2_ in the liver (Figure 6d,e) as well as a depletion of circulating levels of eicosanoids (Figure 4a,b) due to ATP10A deficiency. cPLA_2_ hydrolyzes PC to release AA which can be oxidized to form eicosanoids, bioactive lipids with roles in inflammation and vasculature maintenance (46,47). cPLA_2_ can only act on lipids in the cytosolic leaflet and ATP10A could potentially provide additional PC substrate to this enzyme, derived from the extracellular leaflet or an external source, such as lipoproteins. Interestingly, ATP10A can also translocate GlcCer to potentially promote synthesis of ceramide 1-phosphate (C1P) (Figure 4d), another activator of cPLA_2_ (48). cPLA_2_ hydrolysis of PC also produces LysoPC, a bioactive lipid that may link saturated FAs to insulin resistance (49). Despite the reduction of eicosanoids in the plasma, there was no substantial difference in the levels of circulating LysoPC based on genotype. It is possible that the increased expression of *Pla2g5* (Figure 4c, Supplemental Table 2), a secreted phospholipase, may have compensated for the perturbations in cPLA_2_ signaling and normalized LysoPC levels in the plasma. Further work is needed to better define the specific influence of ATP10A on cPLA_2_ signaling and eicosanoid production.

Prior studies analyzing mice with an overlapping series of radiation-induced deletions encompassing the *p* locus on chromosome 7 implicated *Atp10A* deficiency in diet-induced obesity, insulin resistance indicated by reduced efficacy of insulin in mediating glucose disposal, and hyperlipidemia, and reported that these phenotypes were more severe in females compared to males (26,27). Additionally, the phenotypes were observed in heterozygous mice that inherited the chromosomal deletion maternally, suggesting that the paternal *Atp10A* allele was silenced. In support of this possibility, we found that heterozygous male mice inheriting the *Atp10A* exon 2 deletion maternally displayed a trend toward increased weight gain over the course of the HFD and had significantly shorter bodies and elevated fasting blood glucose compared to mice that inherited the deletion paternally (Supplemental Figure 2a,d,e). However, we found no evidence for increased obesity or metabolic defects in female *Atp10A^+/-^* heterozygous mice inheriting the KO allele paternally or maternally (Supplemental Figure 2), nor did the *Atp10A^-/-^* homozygous mice display alterations to the development of diet-induced obesity or defects in glucose metabolism (Figure 1,2). This difference is most likely because the mice used in previous studies had a large chromosomal deletion that, in addition to *Atp10A*, removed several other genes, some of which are known to be imprinted (50). It is also possible that differences in the strain background or rearing environment alter the susceptibility of *Atp10A* deficient mice to weight gain on a HFD and glucose metabolism perturbations. However, our results are consistent with previous reports in that ATP10A has a stronger influence on lipid metabolism in female mice relative to males (26). We have shown that ATP10A is required to maintain lipid homeostasis and liver insulin sensitivity with HFD feeding, therefore, a therapeutic that acts as an ATP10A agonist could help treat diseases that cause dyslipidemia and hepatic insulin resistance.

## Methods

### Animals

All mouse experiments were approved under the Vanderbilt University Institutional Animal Care and Use Committee. Mice were housed in 12 h light/dark cycles in temperature and humidity-controlled facilities with ad-libitum access to diet and water.

### Creating the mouse model

The *Atp10A* mouse model (knockout allele name: *Atp10Aem1(Vtrg)* (allele designation approval pending)) was created via CRISPR-Cas9 in collaboration with the Vanderbilt Genome Editing Resource. Guide RNAs (crRNA) were created to target *Atp10A* on chromosome 7: Target Intron 1-2: TGACTGCTTAATGATTCGAGG, GAGTGACTGCTAATGATCG, Target Intron 2-3: GGAAAAAGCCCAATTCCACAC, AGCCCAATCCACACAGGAAC. CRISPR editing reagents were delivered into the pronuclei of one-cell fertilized mouse zygotes (C57BL/6J). Approximately 608 bp were deleted using this method: nucleotides 58389679-58390287 (NCBI reference sequence: NC_000073). The resulting pups were biopsied and screened by PCR and Sanger sequencing. The predicted founders were bred to WT C57BL/6J animals and the offspring were genotyped (N1 generation). The offspring with the appropriate genotype were then backcrossed two more times.

### Genotyping

Mice were genotyped using tail DNA. *Atp10A* DNA products were detected via PCR (Q5 DNA Polymerase, NEB) followed by gel electrophoresis; *Atp10A-F* (GTGCACTGTATTTGTCTGCCTGTTCC), *Atp10A-R (*GGTCCTTTGAAGAGATAATGTTCCCAAC).

### Body composition, body length, and tissue mass

WT and experimental mice were fed standard chow or 60% HFD (D12492, Research Diets) ad libitum, starting at the age of 3-12 weeks old (see Supplemental Table 3). Mice were weighed once per week to measure body weight gain over time. On the 12^th^ week on the HFD, body composition was assessed via NMR (LF50 Body Composition Mice Analyzer, Bruker, stock # E140000501). Mice were sacrificed with CO_2_, the body lengths were measured and the mass of the wet tissue was measured using an analytical scale. Tissues were collected and flash frozen when mice were sacrificed.

### *Oral glucose tolerance tests* (OGTT) *and fasting blood glucose/insulin measurements*

Mice were fasted for 5 hours (7AM-12PM) and then an OGTT was performed. Mice were gavaged with 20% w/v dextrose (final 2g/kg body weight), and a tail-vein blood glucose was measured via a glucometer (Accu-Chek, Accu-Chek Aviva Plus Meter) at baseline, 15, 30, 45, 60, 90, and 120 min. The area under of the curve for glucose was calculated using GraphPad Prism. Plasma samples were collected from 5-hr fasted mice via a retroorbital bleed and were used for the insulin assay. Plasma insulin was measured using the Crystal Chem Ultrasensitive Mouse Insulin ELISA Kit (catalog # 90080).

### Plasma lipid and lipoprotein analysis

Plasma was collected, via a retroorbital bleed or cardiac puncture, from 5-hr fasted mice and 5-hr fasted mice that had undergone an OGTT (see Supplemental Table 3). For males and a portion of the female samples: plasma TG and cholesterol were measured using colorimetric kits (note that the TG measurements include free glycerol) (Inifinity, Thermo Scientific, TG catalog #TR22421, chol catalog #TR13421) and plasma FFAs were measured using Abcam’s Free Fatty Acid Assay Kit-Quantification (ab65341). For the rest of the females: plasma samples were measured by the Vanderbilt Hormone Assay and Analytical Services Core, where plasma cholesterol and TG were measured by standard enzymatic assays, and plasma FFAs were analyzed with an enzymatic kit (Fujifilm Healthcare Solutions, catalog #999-34691). Lipoproteins were separated using FPLC on a Superose6 column (GE Healthcare) from 150 μl plasma (single mouse or pooled) and the TG and cholesterol content were measured using colorimetric kits (Inifinity, Thermo Scientific, TG catalog #TR22421, chol catalog #TR13421). AUC was calculated using GraphPad Prism.

### Plasma lipidomics

Plasma was collected, via a retroorbital bleed or cardiac puncture, from 5-hr fasted mice and some samples are pooled plasma (see Supplemental Table 3). Lipid metabolites were extracted from 100 μL plasma by methyl methyltert-butyl ether (MTBE) extraction. The lipid metabolites were then analyzed by HPLC-IM-MS/MS on an Agilent 6560 mass spectrometer using a ZORBAX Extend-C18 RPLC column (Phase A: 0.1% formic acid and 10 mM NH_4_CHOO in water, Phase B: 0.1% formic acid and 10 mM NH_4_CHOO in 60:36:4 isopropanol:acetonitrile: H_2_O) (51). Data alignment and biostatical analysis was performed using Progenesis QI (Waters). Tentative compound identifications were assigned using accurate mass databases and a previously described ion mobility-based filtering method (52).

### RNA Sequencing

Visceral fat was flash frozen in liquid nitrogen and then kept at -80°C until thawed at -20°C in RNA*later* (ThermoFisher, catalog # AM7030). The tissue was then homogenized in QIAzol Lysis Reagent (Qiagen, catalog #79306) using a Bullet Blender (Next Advance, BT24M). The RNA layer acquired after the QIAzol Lysis protocol was cleaned up using the RNeasy Lipid Tissue Mini Kit (Qiagen, catalog #NC9036170). Quality control measurements and Next Generation Sequencing (NGS) was performed on 20 uL of RNA (>10 ng) by Vanderbilt Technologies for Advanced Genomics (VANTAGE); briefly, NEBNext Ultra II Directional RNA kit (Cat no: E7760L) was used and for the sequencing; NovaSeq 6000 and PE150 read lengths were used. Analysis of NGS data was performed by Vanderbilt Technologies for Advanced Genomics Analysis and Research Design (VANGARD).

### Immunoblot analysi

For lysis of liver tissue samples, T-PER (Pierce, Rockford, IL) with phosphatase and protease inhibitors (Sigma) was used. Protein concentration was quantified using BCA Reagent (Pierce, Rockford, IL). Protein extracts (100 µg) were subjected to Western immunoblot analysis. The following primary antibodies were used for detection of: DGAT1 (sc-32861, Santa Cruz Biotech), DGAT2 (sc-66859, Santa Cruz Biotech), phospho-AMPKα^T172^ (#2535, Cell Signaling), AMPKα (#2532, Cell Signaling), phospho-ACC^S79^ (#3661, Cell Signaling), ACC (#3676, Cell Signaling), phospho-IGF-IRβ^Y1131^/IRβ^Y1146^ (#3021, Cell Signaling), phospho-IGF-IRβ^Y1135/1136^/IRβ^Y1150/1151^(#3024, Cell Signaling), IRβ (#3025, Cell Signaling), phospho-IRS-1^S612^ (#3203, Cell Signaling), IRS-1 (#3407, Cell Signaling), phospho-Akt^T308/S473^ (#13038/#4060, Cell Signaling), Akt (#9272, Cell signaling), phospho-GSK-3β^S9^ (#5558, Cell Signaling), GSK-3β (#9315, Cell Signaling), phospho-cPLA_2_α^S505^(#53044, Cell Signaling), cPLA_2_α (#5249, Cell Signaling). Antibody to β-actin (#4970, Cell Signaling) was used to evaluate protein loading in each lane. Immunoblots were developed using the Western Lightning Chemiluminescence Plus detection system (PerkinElmer, Wellesley, MA) according to the manufacturer’s protocol. Intensity of the immunoblot bands was measured using AI600 CCD Imager for chemiluminescent assays (Amersham). Densitometry was performed using ImageJ. For quantification, OD of bands for phosphoprotein was normalized to total protein and β-actin; otherwise, OD of bands for total protein was normalized to β-actin.

### Promethion System

Indirect calorimetry and additional measures of food intake and physical activity were performed in open-circuit indirect calorimetry cages (Promethion System, Sable Systems International) at Vanderbilt Mouse Metabolic Phenotyping Center (MMPC). One week prior to the experiment start date, mice were singly housed for acclimation. On the day of the experiment start date, mice were weighed and body composition was assessed. The mice were placed in the cages of the Promethion system, one mouse per cage. The cages were housed in a light and temperature-controlled chamber. The light cycle was set on a 12:12h cycle (6am-6pm). Mice were left undisturbed for 5 days during which all the measurements were made. The system periodically measured rates of O2 consumption (VO2) and CO2 production (VCO2) for each cage, as well as food intake and physical activity (infrared beam array). Data were processed in time segments (daily 24-hour, and 12-hour dark/12-hour light), and averages calculated for each as well as for the 24-hour period. After 5 days, the mice were removed from the cages, and body composition and weight were measured again. Data was further processed using CalR (www.calrapp.org).

### Measuring liver lipids

Liver FFA, total cholesterol, cholesterol ester, unesterified cholesterol, TG, PLs, and ceramides were measured by the Vanderbilt Hormone Assay and Analytical Services Core using methods described in (53,54).

### Oil Red O

Livers were placed in frozen section medium (Epredia, Neg-50, catalog #22-110-617) and then placed on dry ice. When the frozen section medium was solidified, the livers were placed at -20°C until further processing. Further processing was done by the Vanderbilt Translation Pathology Shared Resource (TPSR): Slides were brought to room temperature and placed in 10% NBF solution for 10 minutes. The Newcomer Supply Tech Oil Red O, Propylene Glycol staining kit (Newcomer Supply, catalog # 12772B) was used for visualization. Slides were then cover slipped with aqueous mounting medium. In regard to Oil Red O scoring: Dr. Katherine Gibson Corley, a board-certified veterinary pathologist at the Vanderbilt TPSR, created the scoring system used to score the Oil Red O stained liver sections. The scoring system is characterized by: 0= no Oil Red O staining, 1=rare and scattered Oil Red O staining, 2=multi-focal and coalescing Oil Red O staining, 3=diffuse Oil Red O staining (in whole tissue). To score the slides, the scorer was blinded to the slide IDs and then scored the slides twice, on two separate days. Then the scores from both days were compared and if they did not match, the scorer viewed the slide again and decided on a final score. The diameter of the Oil Red O positive lipid droplets was measured from slides with a score of 2, using ImageJ. Images of tissues were acquired using light microscopy (Zeiss AxioPlan, Upright Fluorescence Microscope, Germany) with a x5/1.0 or x63/1.0 Oil using the Zeiss Axio Color Camera (Axiocam 208c) and the ZEN 3.1 software (blue edition). Microscope settings were held constant for all experimental groups.

### Statistics

All statistical analysis was done using GraphPad Prism, version 9.5.0 (GraphPad Software). Error bars indicate mean with standard error of the mean (SEM). When more than 2 groups were compared, a 2-way ANOVA was used with Dunnett’s correction for multiple comparisons with a control group. Šidák’s correction was used for comparison of groups of means. Differences between group mean values were tested using a 2-tailed Student’s *t* test or a Mann-Whitney *U* test for nonparametric data. A *P* value of less than 0.05 was considered statistically significant.

### Study approval

The animal protocol was approved by Vanderbilt University Medical Center and IACUC.

### Data Availability

The untargeted lipidomics data is available at the NIH Common Fund’s National Metabolomics Data Repository (NMDR) website, the Metabolomics Workbench (55), where it has been assigned Study ID ST002696. The data can be accessed directly via its ProjectDOI: http://dx.doi.org/10.21228/M83H7N.The reads from the RNASeq data can be found at NCBI (https://www.ncbi.nlm.nih.gov/genbank/samplerecord/) using these accession numbers: SRR24233646 (*Atp10A* WT reads) and SRR24233645 (*Atp10A* KO reads).

## Author contributions

ACN is the first author on this publication; she designed and conducted all of the mouse experiments, collected the tissues analyzed in the paper, and did the microscopy work. Additionally, she created the figures and schematics and wrote the manuscript. EMY conducted all of the western blot experiments, helped create the manuscript figures, and edited the manuscript. LZ helped with the mouse work, tissue collection, and provided intellectual support. BSR conducted the mass spectrometry analysis and analyzed the results. JCM developed the multidimensional mass spectrometry methods and assisted with the lipidomics informatics work. KGC created the Oil Red O scoring system. JAM, JMS and TRG provided funding acquisition, resources, conceptualization, and project administration.

## Acknowledgements

The authors thank the **Vanderbilt Genome Editing Resource** for creating the *Atp10A* mouse lines and assisting with genotyping (supported by the Cancer Center Support Grant (CA68485), the Vanderbilt Diabetes Research and Training Center (DK020593) and the Center for Stem Cell Biology), the **Vanderbilt Division of Animal Care** veterinarians and technicians for their support with mouse husbandry, the **Vanderbilt Metabolic Mouse Phenotyping Center (MMPC)** especially Louise Lantier, for providing the NMR machine for body composition measurements and for performing the indirect calorimetry (DK135073, DK020593, S10RR028101), the **Vanderbilt Analytical services core** for measuring plasma and liver lipids (supported by *NIH grant DK020593 (DRTC))*, the **Translational Pathology Shared Resource (TPSR)** for producing all of the paraffin embedded tissue sections and performing the Oil Red O and CD31 staining (supported by NCI/NIH Cancer Center Support Grant P30CA068485), the **Vanderbilt Cell Imaging Shared Resource (CISR)** for providing the Zeiss LSM880 Airyscan Confocal Microscope used in this study (supported *by NIH grants CA68485, DK20593, DK58404, DK59637, and EY08126;* Zeiss LSM880 Airyscan Confocal Microscope *acquired through NIH* 1 S10 OD021630 1), the **Center for Innovative Technology at Vanderbilt** for providing access to the mass spectrometry instrumentation and lipidomics software, **The Metabolomics Workbench** *(supported by NIH grants U2C-DK119886 and OT2-OD030544),* Sophia Yu, Teri Stevenson, and Staci Bordash for help with mouse experiments, and Bridget Litts for help with finding materials and using equipment. We also thank David Wasserman, Owen McGuiness and David Harrison for helpful discussions. This work was supported by NIH 1R35GM144123 (to TRG), K01AG077038 (to LZ), NIH R01DK109102, R01HL144846, and T35DK7383, The Department of Veterans Affairs BX002223 (to JMS).

## Conflict of Interests

The authors have declared that no conflict of interest exists.

---

**Supplemental Table 1.**
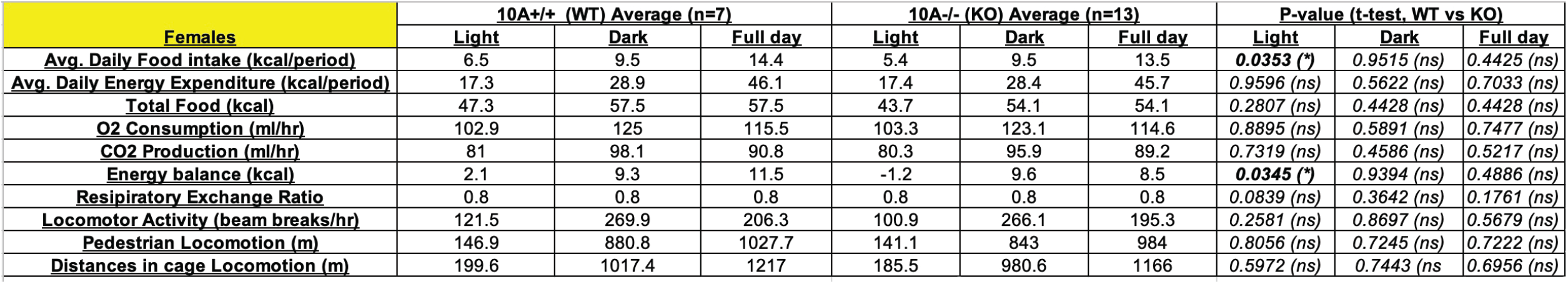
*Atp10A* deletion causes alterations to average daily food intake and energy balance during the light hours in female mice fed a HFD for 13 weeks. The table indicates the parameters measured in the Sable System’s Promethion Indirect Calorimetry System where the mice were single-housed. The values are an average measurement from 5 days in the system with ad libitum feeding. P value by unpaired t-test (*10A^+/+^* n=7, *10A^-/-^* n=13).

**Supplemental Table 2.**
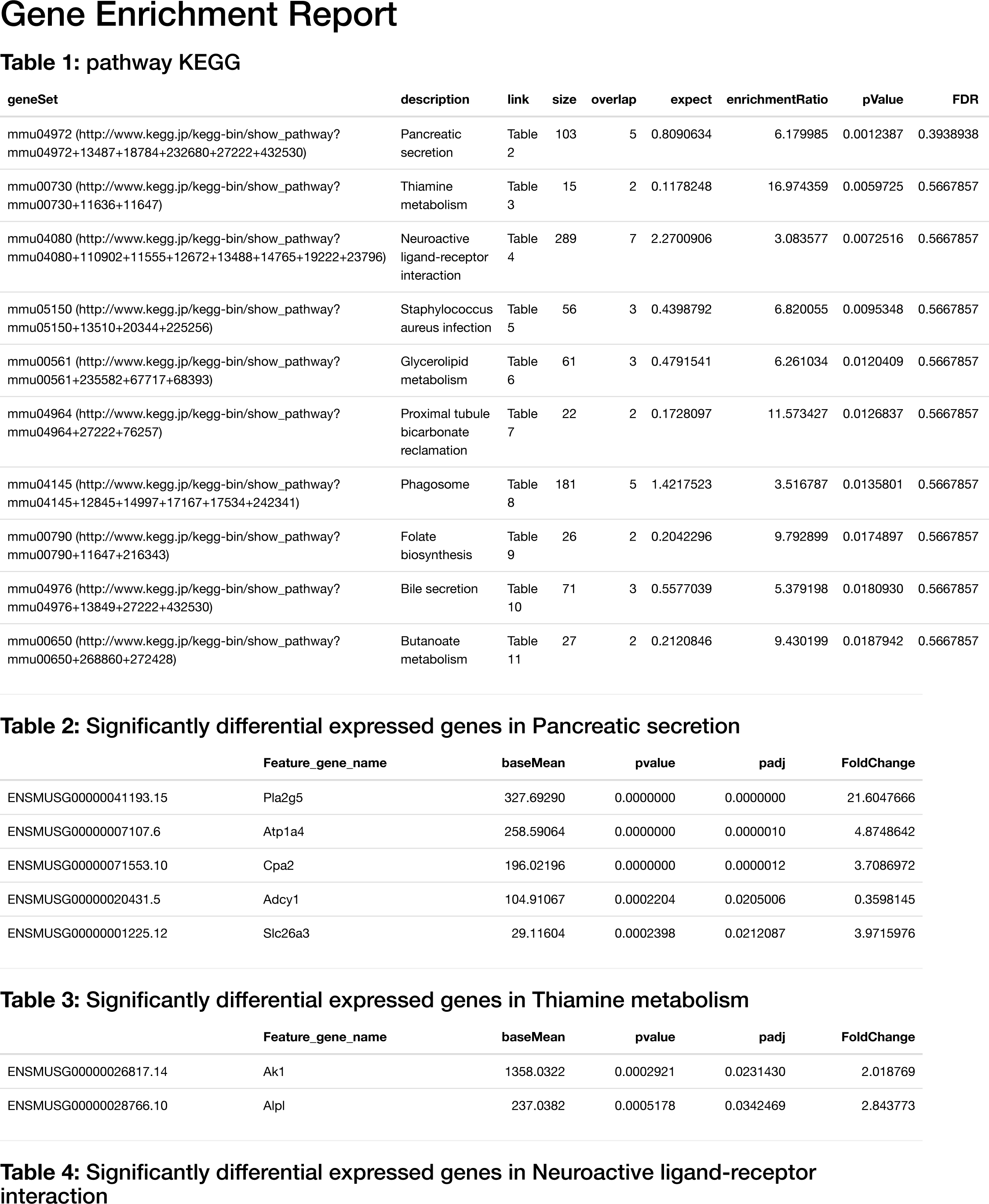
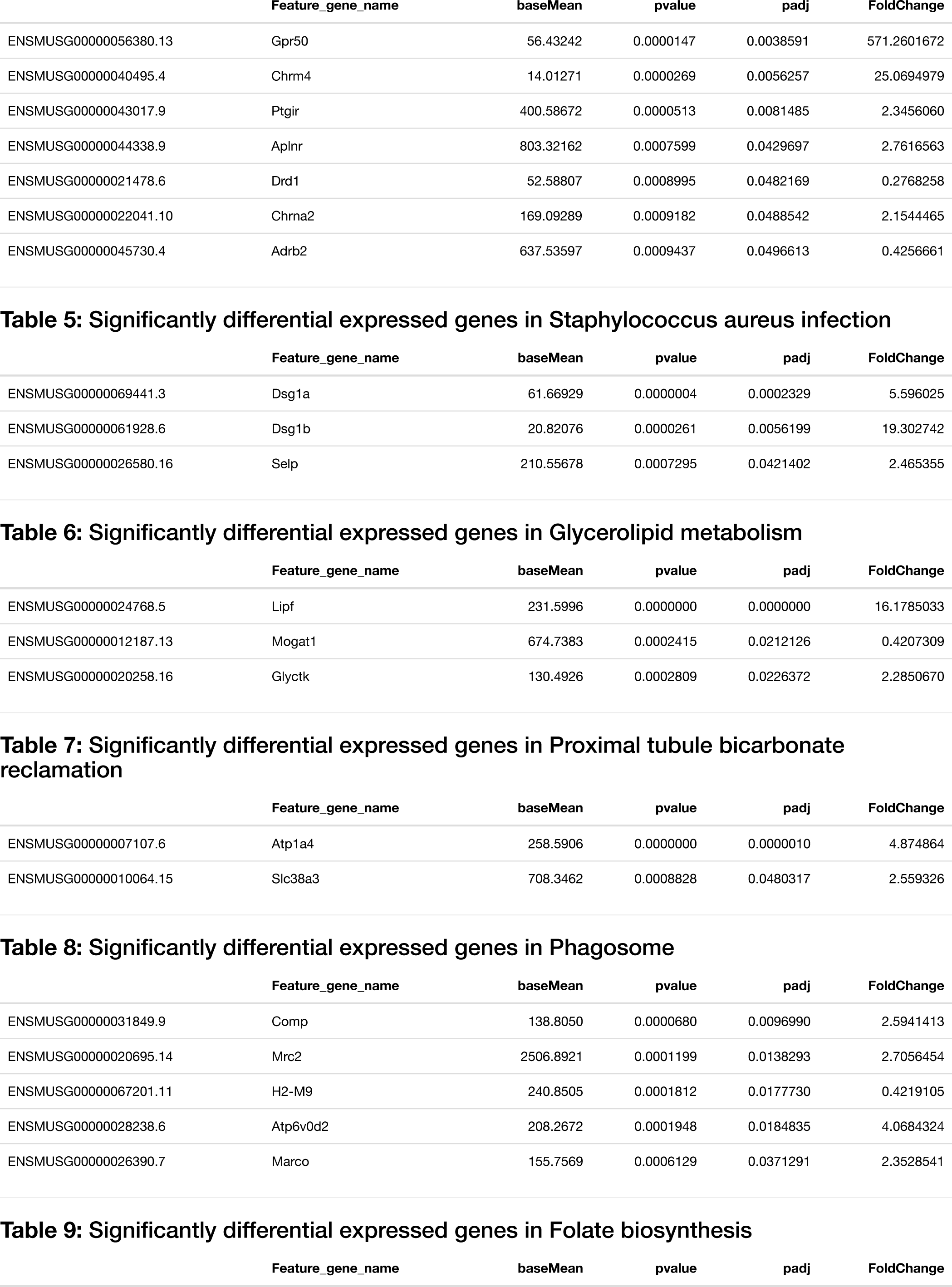
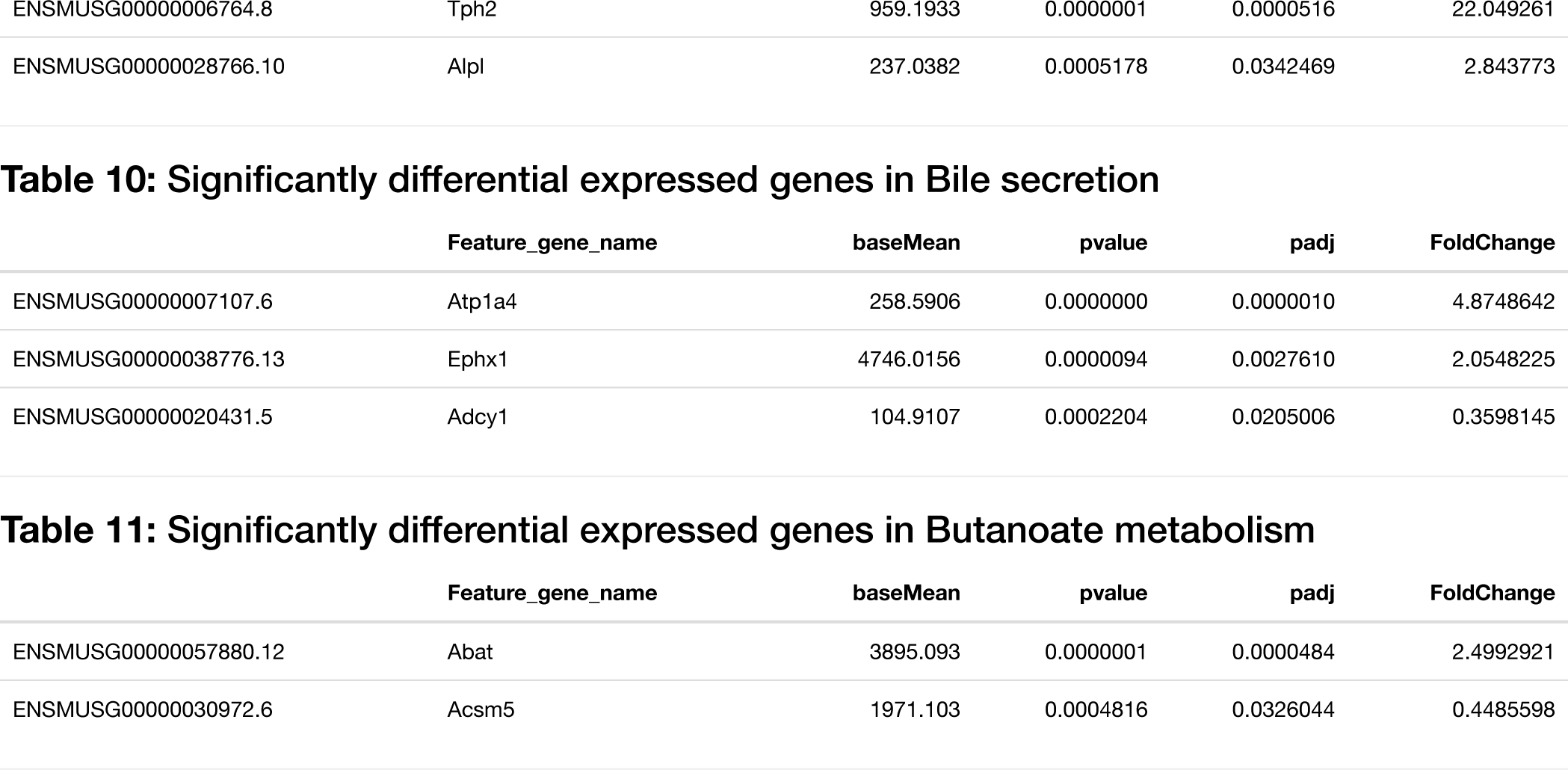
*Atp10A* deletion results in changes to the expression of genes involved in several biological pathways in visceral fat after HFD feeding. Gene enrichment report for visceral fat mRNA gene expression data from *10A*^+/+^ and *10A*^-/-^ female mice (Figure 4C) created using the Kyoto Encyclopedia of Genes and Genomes (KEGG) database. FDR= false discovery rate, padj= p-value adjusted for multiple testing.

Supplemental Table 3. **Characteristics of mice used in this study.** The table includes information on sex, sample size, body mass after HFD feeding, age at time of sacrifice, experimental conditions before sacrifice, and statistical outlier testing.

**Supplemental Figure 1.**
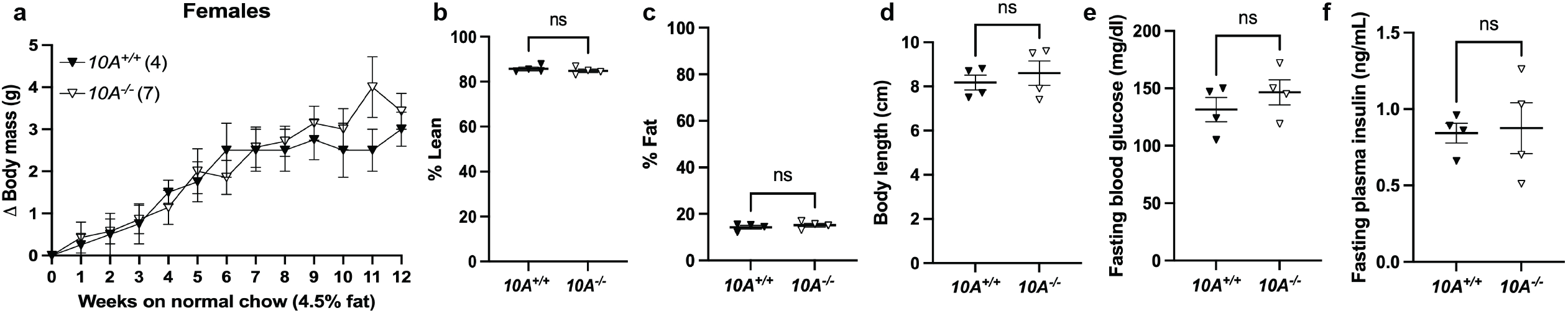
*Atp10A* deletion does not affect weight gain, body composition, or glucose homeostasis in female mice after 12 weeks on normal chow. (a) Weight gain of *10A^+/+^* and *10A^-/-^* female mice over the course of 12 weeks on normal chow (4.5 kcal% fat, Ad lib feeding), (*10A^+/+^* n=4,*10A^-/-^*n=7). (b) Lean and (c) fat body mass was normalized to the combined sum of lean and fat mass to calculate % Lean and % Fat mass (*10A^+/+^* n=4, *10A^-/-^* n=4). (d) Body length of mice was measured after CO_2_ sacrifice (*10A^+/+^* n=4,*10A^-/-^*n=4). (e) Fasting blood glucose was measured a 5 hour fast, via a glucometer (*10A^+/+^* n=4,*10A^-/-^* n=4). (f) Fasting plasma insulin was measured after a 5 hour fast (*10A^+/+^* n=4,*10A^-/-^*n=4). P value by (a) 2-way ANOVA with Sidak’s multiple comparison or (b-f) unpaired t-test.

**Supplemental Figure 2.**
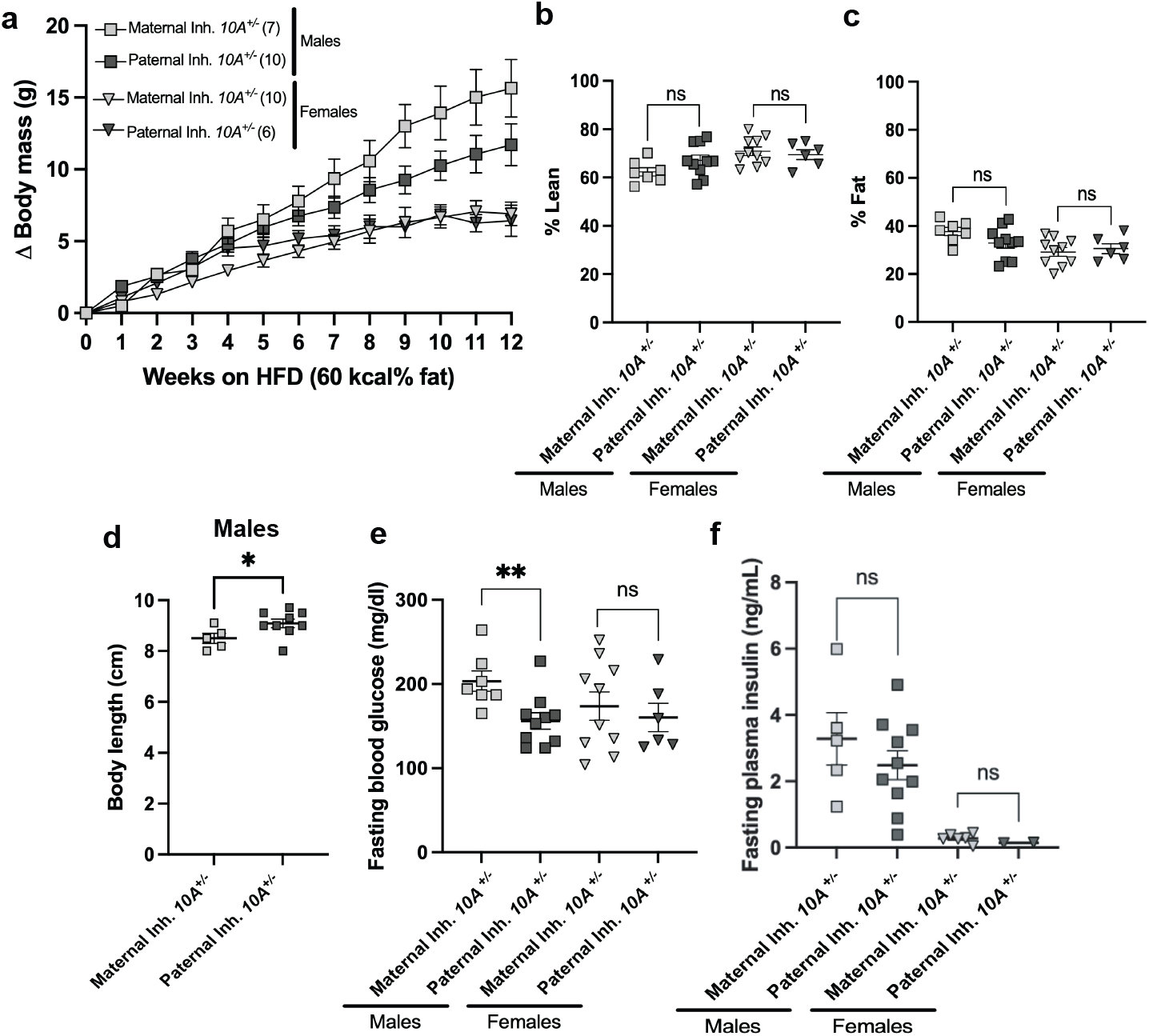
Maternally inherited *Atp10A* deletion leads to smaller bodies and elevated fasting blood glucose in male mice on the 12th week on the HFD. (a) Weight gain of heterozygous males and females inheriting the KO allele maternally (Maternal Inheritance (Inh.), *10A^+/-^* dam X *10A*^+/+^ sire) or paternally (Paternal Inh, *10A*^+/+^ dam x *10A^+/-^* sire) over the course of 12 weeks on a HFD (60 kcal% fat, Ad lib feeding) (**Males**: Maternal Inh.*10A^+/-^*n=7, Paternal Inh.*10A^+/-^* n=10; **Females**: Maternal Inh.*10A^+/-^* n=10, Paternal Inh.*10A^+/-^* n=6). (b) Lean and (c) fat body mass were normalized to the combined sum of lean and fat mass to calculate % Lean and % Fat mass (**Male**: Maternal Inh.*10A^+/-^* n=7, Paternal Inh.*10A^+/-^* n=10; **Female**: Maternal Inh.*10A^+/-^* n=10, Paternal Inh.*10A^+/-^* n=6). (d) Body length of male mice was measured after CO_2_ sacrifice, *P=0.0490. (Maternal Inh.*10A^+/-^*n=4, Paternal Inh.*10A^+/-^* n=9). (e) Fasting blood glucose was measured after a 5 hour fast, via a glucometer, **P=0.0081. (**Male**: Maternal Inh.*10A^+/-^*n=7, Paternal Inh.*10A^+/-^* n=10; **Female**: Maternal Inh.*10A^+/-^* n=10, Paternal Inh.*10A^+/-^*n=6). (f) Fasting plasma insulin was measured after a 5 hour fast (**Male**: Maternal Inh.*10A^+/-^* n=6, Paternal Inh.*10A^+/-^* n=10; **Female**: Maternal Inh.*10A^+/-^*n=5, Paternal Inh.*10A^+/-^* n=2). (a) P value by 2-way ANOVA with Sidak’s multiple comparison or (b-f) unpaired t-test.

**Supplemental Figure 3.**
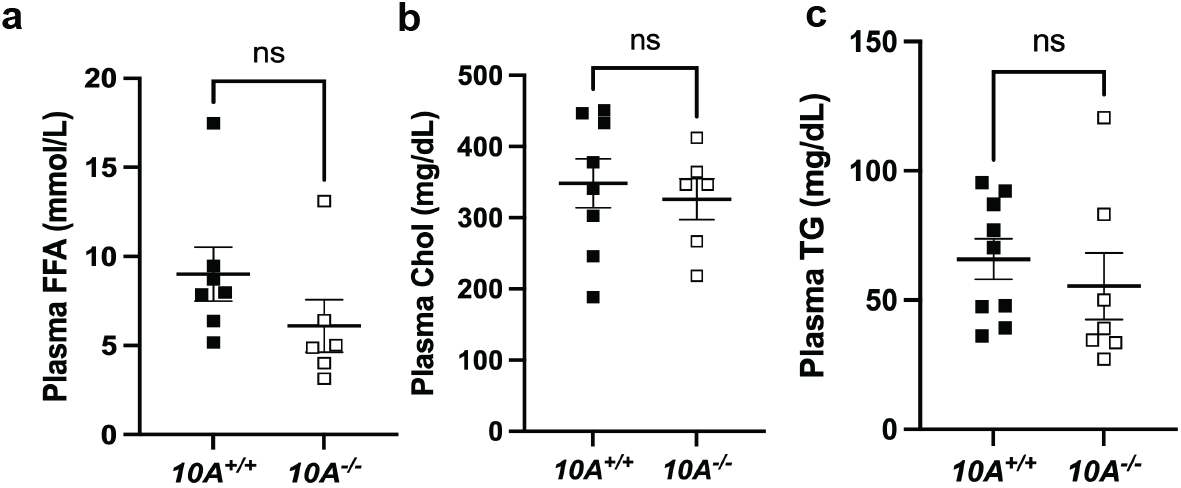
*Atp10A* deficiency does not result in dyslipidemia in male mice after HFD feeding. (a-c) Free fatty acids (FFA), cholesterol (chol), and triglycerides (TG) were measured in plasma from males after a 5 hour fast. P value by unpaired t-test. (a, *10A^+/+^* n=7, *10A^-/-^* n=6; b, *10A^+/+^* n=8, *10A^-/-^* n=6; c, *10A^+/+^* n=9, *10A^-/-^* n=7).

**Supplemental Figure 4.**
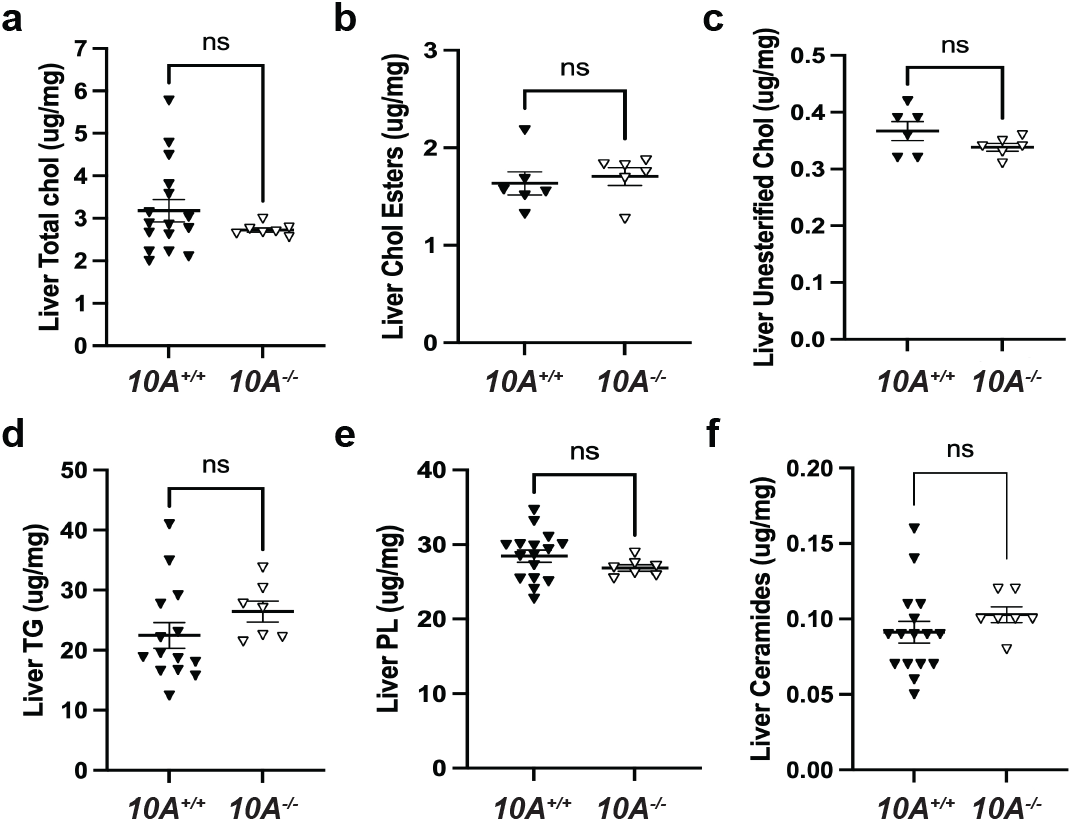
*Atp10A* deletion does not affect total amounts of liver cholesterol, TGs, PLs, or ceramides in female mice after 12 weeks of HFD. Total (a) cholesterol, (b) cholesterol esters, (c) unesterified cholesterol, (d) TGs, (e) PLs, and (f) ceramides were measured from flash frozen livers via gas chromatography. Livers were collected after a 5 hour fast or after a 5 hour fast followed by an OGTT. P value by unpaired t-test (Total cholesterol: *10A^+/+^* n=16, *10A^-/-^* n=7; Cholesterol esters: *10A^+/+^* n=6,*10A^-/-^* n=6; Unesterified cholesterol: *10A^+/+^* n=6,*10A^-/-^* n=6; TG: *10A^+/+^* n=14, *10A^-/-^* n=7; PLs: *10A^+/+^*n=16, *10A^-/-^* n=7; Ceramides: *10A^+/+^* n=16, *10A^-/-^* n=7).

## Notes

http://dx.doi.org/10.21228/M83H7N

### Competing Interest Statement

The authors have declared no competing interest.

### Summary of Updates

Reviewers of the original manuscript did not like having both major phenotypes (dyslipidemia and endothelial dusfunction) presented in the same manuscript without clear mechanistic links between them. In response, we are splitting the work into two manuscripts and the first new manuscript on dyslipidemia has the most similar title to the original and is being used to substitute for the original submission. We will submit the second new manuscript to biorxiv in a few months.

